# Modulation of the Biophysical and Biochemical Properties of Collagen by Glycation for Tissue Engineering Applications

**DOI:** 10.1101/2022.03.03.482886

**Authors:** Mina Vaez, Meisam Asgari, Liisa Hirvonen, Gorkem Bakir, Sebastian Aguayo, Christina M. Schuh, Kathleen Gough, Laurent Bozec

## Abstract

The structural and functional properties of collagen are modulated by the presence of intramolecular and intermolecular crosslinks. Advanced Glycation End-products (AGEs) can produce intermolecular crosslinks by bonding the free amino groups of neighboring proteins. In this research, the following hypothesis is explored: The accumulation of AGEs in collagen decreases its proteolytic degradation rates while increasing its stiffness. Fluorescence Lifetime Imaging (FLIM) and Fourier-transform infrared spectroscopy (FTIR) detect biochemical changes in collagen scaffolds during the glycation process. The accumulation of AGEs increases exponentially in the collagen scaffolds as a function of Methylglyoxal (MGO) concentration by performing autofluorescence measurement and competitive ELISA. Glycated scaffolds absorb water at a much higher rate confirming the direct affinity between AGEs and interstitial water within collagen fibrils. In addition, the topology of collagen fibrils as observed by Atomic Force Microscopy (AFM) is a lot more defined following glycation. The elastic modulus of collagen fibrils decreases as a function of glycation, whereas the elastic modulus of collagen scaffolds increases. Finally, the enzymatic degradation of collagen by bacterial collagenase shows a sigmoidal pattern with a much slower degradation rate in the glycated scaffolds. This study identifies unique variations in the properties of collagen following accumulation of AGEs.

## Introduction

Collagen type I is the most abundant extracellular matrix (ECM) protein that is responsible for conferring mechanical resilience to connective tissues.^[1]^ To achieve this, both collagen ultrastructure and biochemistry present anatomical specific variations. These variations ensure that collagen-rich connective tissues can accommodate the different environmental demands defined by the functional properties of the tissue as established by Wolff’s Law.^[2]^ Thus, the fine tuning of collagen properties is critical for continued tissue homeostasis. Any significant disruption of these properties is often associated with pathological conditions such as fibrosis ^[3]^, rheumatoid arthritis ^[4]^, and cancer.^[5]^

Tissue engineering has relied on collagen as a native structural protein to engineer scaffolds and, membranes with significant success over the last three decades.^[6]^ The biocompatibility, accessible-chemical functionalization, and in vivo turn-over of collagen are undeniable assets for collagen scaffolds and membranes to be developed towards clinical applications.^[7–8]^ Yet, the most significant limitations of engineered collagen scaffolds are their poor mechanical property, poor structural stability, and rapid degradation. The fine tuning of in vitro collagen scaffold properties is not yet on par with that of in vivo tissue properties. Chemical and physical cross-linking methods have been used to control the mechanical and biological stability of reconstituted collagen assemblies.^[9]^ The most common chemical cross-linking reagents are glutaraldehyde (GTA), hexamethylene diisocyanate (HMDI), and 1-ethyl-3-(3-dimethylaminopropyl) carbodiimide (EDC).^[10–11]^ Photoreactive agents (e.g., riboflavin) and plant extracts (e.g., genipin) have also been used.^[12–13]^ These chemical exogenous collagen crosslinking methods are associated with cytotoxicity, calcification, and foreign body response, which usually overshadow their cross-linking potential.^[14]^ The use of physical approaches such as dehydrothermal (DHT) and UV irradiation has also been evaluated to avoid the cytotoxicity associated with the chemical cross-linkers.^[15–16]^ However, physical crosslinking methods include heating, drying, and irradiation and cannot yield sufficient crosslinking degrees.^[17–18]^ Therefore, there is a need for cross-linking agents that are optimal in low toxicity, and that have the ability of conferring mechanical advantages while not adversely affecting long-term tissue homeostasis. To preserve as much of the composition and structure of the ECM as possible and to mimic the collagen properties as found in tissues, a possible solution would be to selectively re-engineer collagen native crosslinks in new scaffolds.

Under physiological conditions, collagen undergoes natural cross-linking via the enzymatic lysyl oxidase and transglutaminase pathways, as well as nonenzymatic glycation. Initially, collagen fibrils are stabilized by intra- and intermolecular crosslinks that are catalyzed by enzymes of the lysyl oxidase and transglutaminase families. However upon maturation and aging the proportion of enzymatic-mediated crosslinks decreases and the proportion of glycation-mediated crosslinks increases.^[19]^

Glycation is the reaction of carbonyl groups of reducing sugars with free amino groups of lipids and proteins to form a Schiff base, which then undergoes a time-dependent rearrangement to form a fairly stable Amadori product.^[20]^ These structures are still reactive and convert to stable substances called advanced glycation end-products (AGEs). AGEs formation results from long-time exposure of proteins to reducing sugars since glycation of collagen is a process without catalysis. The low turnover of collagen causes AGEs to accumulate within the collagen fibrils in our tissues and organs during normal aging or some pathological conditions such as Alzheimer’s disease ^[21]^, and diabetes. In diabetic conditions, glycation is expected to proceed faster due to an increase in available free sugars that are available to react with collagen residues.^[22]^

AGE-mediated crosslinks are known to alter the physical characteristics and thermal denaturation of collagen structures. A previous study showed that glucosepane (the most abundant and relevant AGE-mediated crosslink) is associated with an increased denaturation temperature, reduced density of collagen packing, and increased porosity to water molecules.^[23]^ Gautieri et al. showed that AGEs reduce tissue viscoelasticity by severely limiting fiber–fiber and fibril–fibril sliding and brittle failure mode in tendons treated with <u>M</u>ethyl<u>g</u>ly<u>o</u>xal (MGO).^[24–25]^ Glycation not only results in a modification of the physical properties of the collagen but also modifies collagen interaction with key molecules like enzymes (e.g. collagenase) that lead to enzyme resistance.^[26]^

Although the effect of glycation on collagen tissue properties has been investigated mostly at the tissue level, there is an evident lack of knowledge in the basic science literature explaining the biomechanical impact of AGE-mediated crosslinks on the functional and structural properties of collagen at both the nanoscale (single fibrils) and mesoscale (bundles of fibrils). Thus, in the current work, we investigate the effects of MGO induced AGE-crosslinks on collagen structural ordering, stiffness, water sorption, and, enzymatic degradability by combining multi-scale imaging and mechanical testing via Atomic Force Microscopy (AFM), Attenuated Total Reflectance-Fourier Transform Infrared Spectroscopy (ATR-FTIR), and time-lapse digital imaging. This study identifies unique variations in the properties of collagen following in vitro tissue glycation by MGO and proposes this method of collagen crosslinking as a means to modulate the biophysical properties of collagen fibrils and scaffolds prior to cell seeding or clinical implantation.

## Results and discussion

### ● In vitro engineering of collagen

In connective tissues, the structural and functional properties of collagen are modulated by the presence of intramolecular and intermolecular crosslinks. These crosslinks vary in types and amount depending on the tissue site, associated pathology, or aging.^[26, 30–31]^ Understanding the relation between collagen properties and its crosslinking has been the subject of numerous research over the last 50 years. The work by Bailey et al. paved the way towards using advanced analytical techniques to define and measure the precise amount and type of crosslinks present in a tissue.^[32]^ However, isolating a single crosslink subtype and understanding its impact on the collagen properties remains challenging. While several groups have added exogeneous crosslinks to a tissue biopsy in vitro ^[33–34]^, it is difficult to decouple the composite response from the pre-existing and newly added crosslinks on the properties of collagen. To elucidate the impact of newly formed crosslinks such as those generated from glycation, we propose to use a tissue engineering approach to create collagen fibrils and scaffolds that are structurally reminiscent of native collagen (as found in tissue) without pre-existing crosslinks. These collagen scaffolds have been used widely in clinical applications, including skin ^[35]^, muscle ^[36]^, tendon ^[37]^, cornea ^[38]^, and bone ^[39]^. Figure 1b and Figure 1c present AFM images of a collagen fibril and scaffold that both have been engineered using this approach from a solution of neutralized single collagen molecules (from rat tails) and compared to native rat tail tendons from which the single collagen molecules are extracted (Figure 1d). In vitro collagen fibrils’ length and diameter, vary depending on growth conditions (pH, temperature, and ionic strength) and are typically smaller (in diameter) than those found in tissue, as shown in Figure 1e and Figure 1f (collagen scaffold fibril diameter: 126.1±0.8 nm, rat tail fibril diameter: 269.2±5.6 nm). Based on the results of a turbidity test, it has been determined that the self-assembly of collagen occurs in three phases; (a) lag phase, in which collagen fiber precursors, e.g., dimers and trimers form and turbidity does not change, (b) growth phase where collagen fibrils are formed; the collagen solution becomes turbid and the turbidity increases with time, and (c) a plateau phase in which turbidity stops increasing.^[40]^ In vitro, this fibrillogenesis results in the formation of stable collagen fibrils in which the collagen monomers are held together by weak hydrogen interactions as there are no mechanisms ab-initio to generate covalent crosslinks in between these collagen monomers. From a tissue engineering approach, the two most important structural parameters for collagen fibrils are the presence of the characteristic D-banding periodicity along the long axis of the fibrils combined with a homogeneous cylindrical aspect of the fibrils.^[41–42]^ Under appropriate conditions, collagen molecules self assemble to form microscopic fibrils, fibril bundles, and macroscopic fibers that exhibit D-banding periodicity indistinguishable from native collagen fibers.^[41]^ These features can be readily observed on all the samples’ images in Figure 1. The presence of D-banding on the fibrils acts as structural markers for intact and native collagen as found in native tissues.^[43]^ There have been several studies investigating the link between variation in the D-banding periodicity and pathology or tissue damage. For example, collagen fibrils with diminished or nearly absent D-periodic banding and irregular cross-sectional profiles of collagen fibrils have been found in the dermal tissues of patients with diseases like Ehlers-Danlos Syndrome (EDS IV) and spontaneous coronary artery dissection (sCAD).^[44–47]^ Localized variation in the shape of the D-banding was reported in some abnormalities like arthrogryposis, renal dysfunction and cholestasis (ACR) syndrome.^[48]^ Finally, local mechanical stress changes have been associated with variations in D-banding periodicity ^[49]^ reinforcing the requirement for tissue engineering approaches to ensure the presence of D-banding periodicity of the fibrils when preparing collagen scaffolds.

**Figure 1.**
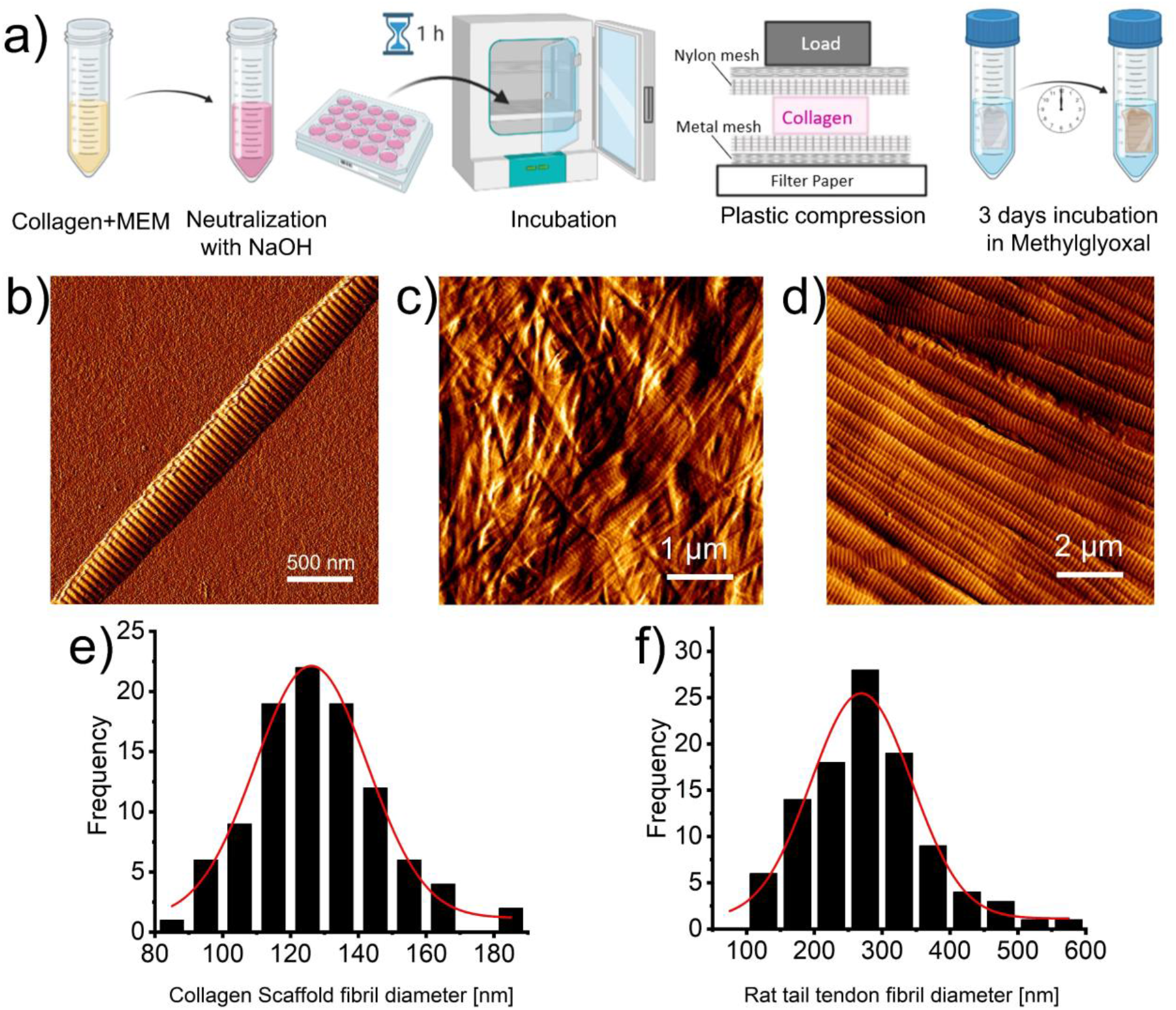
In vitro engineering of collagen. a) Schematic representation of the step-by-step engineering of plastically compressed collagen scaffolds. AFM topological images of b) single collagen fibril (source rat tails) c) collagen fibrils in scaffold (source rat tails), d) extracted collagen fibrils in rat tail tendon. All collagen samples were imaged mounted on a glass slide and imaged in ambient conditions in contact mode. e) Distribution of the measured collagen scaffold fibril diameter: Median (±SD) = 126.1±0.8 nm (n=100); f) Distribution of the measured rat tail fibril diameter: Median (±SD) = 269.2±5.6 nm (n=100).

### ● Modulating collagen glycation

Among the reducing sugars, the most reactive sugar in the body is MGO (50,000-fold more reactive than glucose [21]), which makes it an important glycating agent despite the overall low MGO concentration found in tissue. MGO is formed as a by-product of glycolysis, degradation of glycated proteins, and lipid peroxidation and, under physiological circumstances, detoxified by the glutathione-dependent glyoxalase defense system [18, 19]. Therefore, in aging and diabetes, where the glutathione level is decreased in tissues, MGO concentration would increase. In general, MGO is predominantly associated with diabetes ^[50]^ and its associated pathologies. However, the extensive review by Talukdar et al. in 2009, concluded that it has not been proven that MGO by itself significantly contributes to the suggested deleterious effects on the host. However, MGO can provide both antimicrobial and anti-cancer effects ^[51]^ to affected tissues. In humans, the MGO levels have been determined in plasma and in tissues of healthy individuals at ~60-250 nM and ~1–5 μM, respectively.^[52–53]^ MGO forms AGE residues in proteins largely on arginine residues and, to a much lesser extent, lysine. The major MGO-derived AGE is MGO-derived hydroimidazolone 1 (MG-H1) (90% of all MGO adducts) resulting from the reaction between MGO and arginine amino acids. Some of lysine-derived AGEs are 1,3-di(*N*-lysino)-4-methyl-imidazolium (MOLD), *N*-carboxymethyl-lysine (CML), and *N_ε_*-(1-carboxyethyl)lysine (CEL). Also, Methylglyoxal-Derived Imidazolium Crosslink (MODIC) is an arginine-lysine-derived AGE ^[54]^.

Since MGO is one of the major glycation agents in vivo and rapidly induces the formation of AGEs in vitro (in a manner of hours and days), we functionalized the scaffolds with MGO by incubating them in PBS solution, containing 10 mM, 25 mM, 50 mM, and 100 mM MGO for 3 days (corresponding to a 4 to 6-fold increase when compared to the native concentration of MGO in humans). To detect the formation of MGO-derived crosslinks directly on the collagen, we performed ATR-FTIR. In Figure 2a, microATR-FTIR spectra of non-glycated and glycated with 100 mM MGO collagen scaffolds are displayed and compared. The carbohydrate double band peaking at 1080 cm^−1^ and 1031 cm^−1^ has a larger amplitude in the glycated scaffold spectrum with respect to the control one (non-glycated collagen scaffold). This enhanced intensity in the carbohydrate double band (sugar) is associated with the accumulation of the glycation products in the MGO-treated sample but is also found not to vary (in intensity) as a function of the concentration of MGO (data not shown). To further confirm the formation of MGO-derived AGEs into the scaffolds, we measured the autofluorescence of the glycated collagen scaffolds. As presented in Figure 2b, glycated scaffolds showed an exponential time-dependent and dose-dependant increase in AGE-associated fluorescence (340/420 nm) when compared to the control sample (non-glycated). For example, 100 mM MGO sample exhibited ca. five-time higher fluorescence signal after 3 days incubation compared to the control (Control: 4031±284 vs. 100 mM MGO: 19997±1549 RFU). It is also interesting to note that the control sample increases its autofluorescence over time. This is likely related to residual AGE sidechains fragments present around the rat tails collagen molecules as reported elsewhere.^[20, 55]^ The fluorescence of the glycated scaffolds tends to plateau at 72h suggesting that the glycation reaction may have been completed (or slowed down) by that time. It is, in fact, anticipated that MGO-derived crosslinks form in a few hours up to a few days.^[20]^ The results found in our approach correlate with other studies, demonstrating an increase in collagen autofluorescence as those reported in collagen gel incubation with glucose-6-phosphate(G6P) ^[55]^ and tendon of diabetic animals.^[56]^

**Figure 2.**
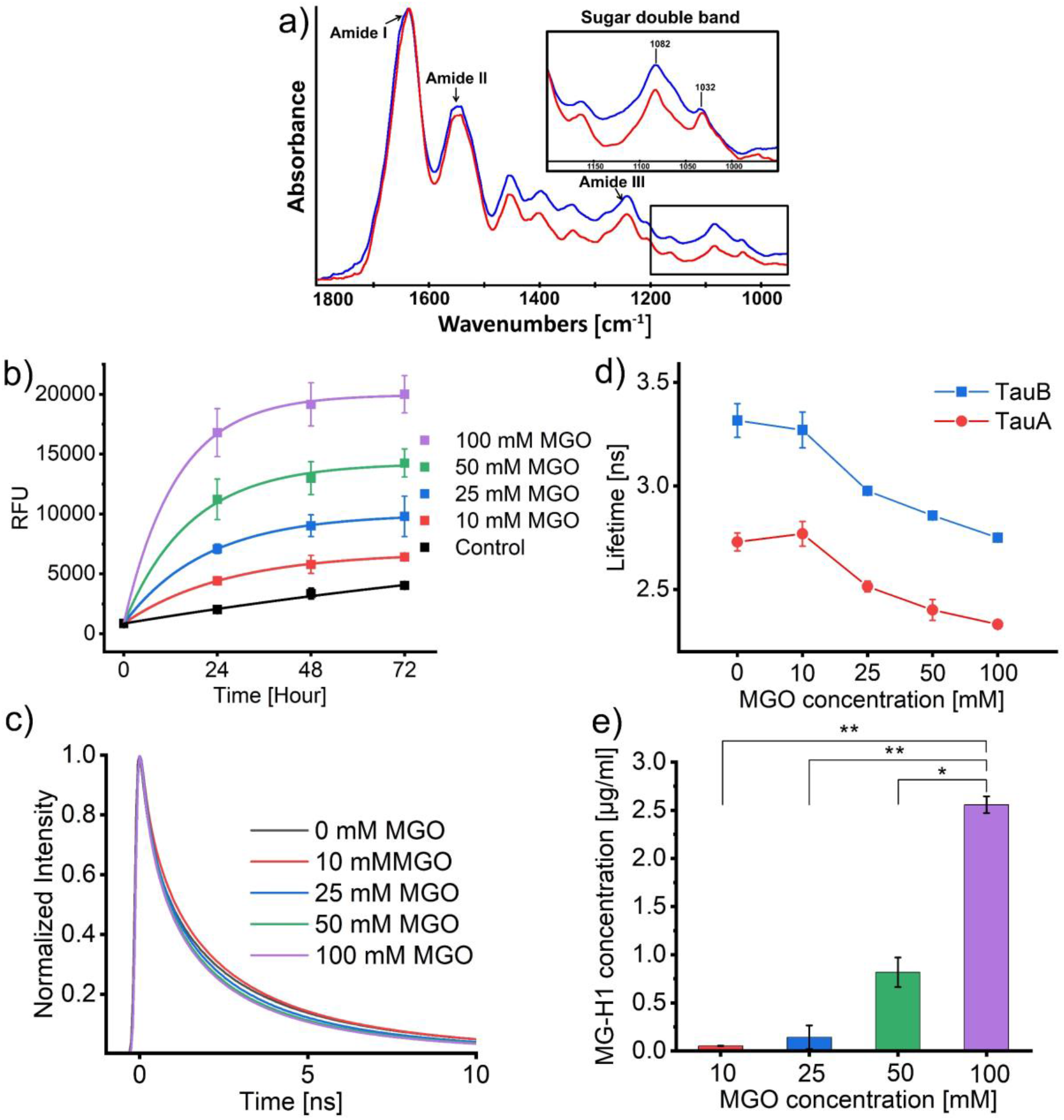
Confirmation of AGE-mediated crosslinking formation. a) Representative infrared spectra of non-glycated (red) and glycated with 100 mM MGO (blue) collagen scaffolds spectra obtained from micro-ATR-FTIR images (not shown) and presenting an enhancement of the carbohydrate double band (sugar) intensity (1032 & 1082 cm^−1^) in the glycated sample; b) Time-dependent autofluorescence profiles of control and glycated collagen scaffolds over a 3-day incubation period. Time point data are presented as Mean autofluorescence measured ± SD (n=3); c) Representative fluorescence lifetime decays of control and glycated collagen scaffolds (zoomed into the first 10 ns of the decay); d) Mean (± SD) fluorescence lifetimes of collagen scaffolds (TauA: average lifetime calculated from the two fastest fitted decay components; TauB: average lifetime calculated from all three fitted decay components) (n=6). e) Histograms presenting levels of MGO-derived MG-H1 (Mean (± SD)) quantified by competitive ELISA (n=3). Statistical significance **P<0.01, *p<0.05 defined using One-way ANOVA with Tukey-test.

In parallel, we explored how the fluorescence lifetime of the glycated collagen decreased as a function of the MGO concentration increase. Time-resolved fluorescence profiles of glycated collagen scaffolds indicated that the fluorescence intensity decayed faster (shorter lifetime) with increasing MGO concentration, as shown in Figure 2c. The mean fluorescence lifetimes, as presented in Figure 2d, were calculated subsequently using equation (1) and equation (2). We found that as the MGO concentration increased, both *τ_A_* and *τ_B_* values decreased from 2.73 ns (control) to 2.33 ns (100mM MGO) for *τ_A_* and from 3.32 ns (control) to 2.75 ns (100 mM MGO) for *τ_B_*. Since the fluorescence lifetime is an intrinsic property of a fluorophore and is not influenced by fluorophore concentration ^[57]^, the reduction in fluorescence lifetime of different glycated scaffolds can be explained through the formation of different fluorescent species or AGEs in our case. This decrease in lifetime values demonstrates that new crosslink species formation is dependent on the MGO concentration. In a similar approach, Fukushima ^[58]^ measured both *τ_A_* and *τ_B_* values in ribose-glycated collagen scaffolds. Interestingly, they found that *τ_A_* did not decrease as a function of glycation and suggested that the decrease in *τ_B_* was associated with the formation of AGEs. The values of *τ_B_* and its trend measured by Fukushima et al., as a function of glycation is in good agreement with the values found in our study. However, the values of *τ_A_* found in our study are higher than that reported by Fukushima et al. This may be explained by the source of collagen used in our respective experiments or by the species generated from MGO when compared to ribose.

However, not all MGO-derived AGEs are fluorescent, and measuring the presence of glycation end-products by autofluorescence and FLIM measurements underestimates the amount or types of AGEs present in collagen. The MGO-derived AGEs can be classified into either a fluorescent or nonfluorescent group based on their ability to emit fluorescence. For example, argpyrimidine is an AGE by-product belonging to the fluorescent AGEs group while imidazolones are nonfluorescent AGEs ^[54]^. To further quantify the formation of MGO-derived AGEs, we employed competitive ELISA for MG-H1. As presented in Figure 2e, the assay showed a dose-dependent exponential increase in the MGO-derived AGEs formation of glycated scaffolds. 100 mM MGO produced 2.56 μg/ml MG-H1 in 100 μg/ml collagen scaffold samples. Since our competitive ELISA measurement targeted the MGO-derived hydroimidazolone 1 (MG-H1) which accounts for 90% of all MGO adducts, we can confirm that our in vitro glycation process can be used to mimic native glycation occurring in the human body despite using MGO solely as the glycation reaction initiator.

### ● Water sorption in glycated collagen (H_2_O/D_2_O exchange)

It has been demonstrated that AGEs have a hydrophilic attraction with the interstitial water present within the collagen fibrils.^[59]^ While tightly bound water molecules are known to stabilize the triple helix by participating in the H-bond backbone, interstitial water is responsible for maintaining the mechanical stability of collagen fibrils. A reduction or absence of interstitial water leads to an increase in the brittle behavior of the collagen fibrils.^[60]^ In collagen, AGEs contain hydrogen bond donors and acceptors that bind to water molecules, and this interaction is thought to be one of the mechanisms to withhold the water within the fibrils. We explored the phenomenon of water sorption in our collagen scaffolds as a function of the MGO glycation by using H_2_O/D_2_O exchange.

To screen the hydrophilic interactions between the AGEs formed and interstitial water (present in the collagen fibrils) by deuteration, we immersed our glycated samples in D_2_O for 48 hours prior to recording their infrared spectra. The vibrational bands of liquid H_2_O are 3530 cm^−1^, 3200 cm^−1^, and 1645 cm^−1^. The vibrational bands of liquid D_2_O are 2625 cm^−1^, 2418 cm^−1^ and 1210 cm^−1^.^[61]^ Spectra recorded after 48hr exposure of scaffolds to D_2_O is shown in Figure 3a. All the spectra have been normalized to the amplitude of the amide I band that allows a direct comparison between the spectra recorded in the presence of H_2_O and D_2_O ^[62]^. The H_2_O/D_2_O exchange spectra indicate that during the soaking period D_2_O naturally displaced H_2_O as demonstrated by the very strong D_2_O absorbance band observed at 2418 cm^−1^. As soon as the D_2_O saturated collagen sample had been mounted on the ATR crystal, the D_2_O started evaporating (in ambient conditions) and the H_2_O present in the ambient air was absorbed by the collagen sample. This was confirmed by recording a rapid decrease in the intensity of D_2_O absorbance at 2418 cm^−1^ while the intensity of H_2_O band at 3200 cm^−1^ increases over the same period.

**Figure 3.**
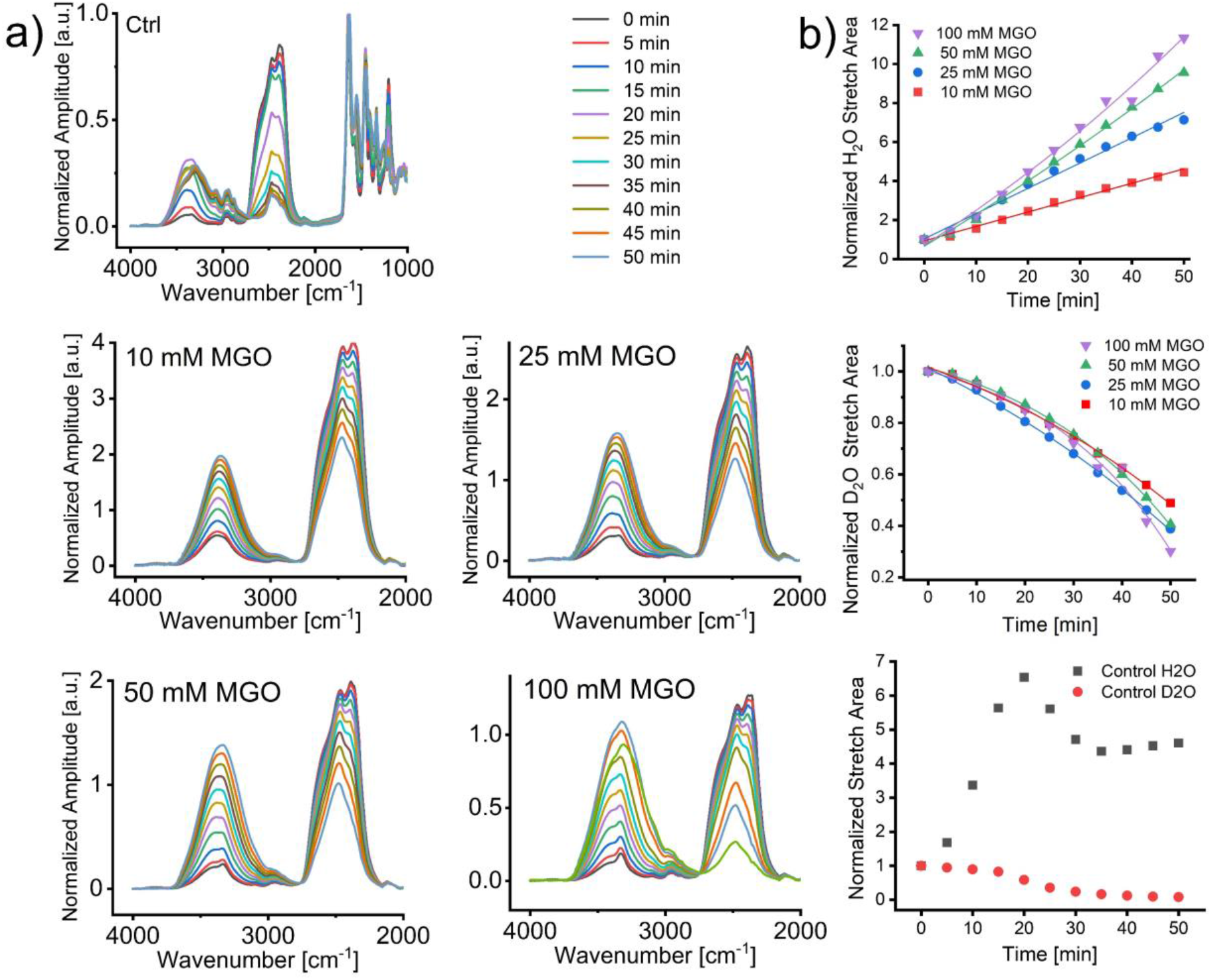
H_2_O/D_2_O exchange in glycated collagen. a) Representative time series of ATR infrared spectra from control and glycated collagen scaffolds mounted directly on the ATR window of the FTIR spectrometer and undergoing D_2_O/H_2_O exchange for 50 min. Spectra were normalized to the Amide I and plotted using the resultant normalized amplitude in arbitrary unit of absorbance; b) Graphical representation of the time-dependent rate of H_2_O sorption and D_2_O evaporation for glycated and control collagen scaffolds. Data points represent the area below the H_2_O and D_2_O stretch bands respectively. For H_2_O sorption and D_2_O evaporation, the data profiles were fitted with exponential growth and decay fits respectively (R^2^>0.95).

The rates at which D_2_O is replaced by atmospheric H_2_O can be measured by monitoring the area of the residual intensity of H_2_O absorbance peak near 3200 cm^−1^ and D_2_O absorbance peak near 2400 cm^−1^ as a function of time. Rates of H_2_O sorption and D_2_O evaporation are presented in Table 1. As shown in figure 3b the rate of D_2_O evaporation remained independent of the level of glycation present in each scaffold. This result confirmed that there are no chemically favorable mechanisms to withhold D_2_O within the collagen scaffold (glycated or not) as D_2_O does not interact with the hydrophilic groups engineered through the glycation process. However, the rate of water sorption varied accordingly to the level of glycation present in the collagen scaffold as measured by the Normalized Peak Area (NPA) over time. As such, the water sorption rate increased from (0.074 ±0.002 NPA.min^−1^) for the 10 mM MGO glycated scaffold to (0.2102+/-0.007) NPA.min^−1^ for the 100 mM MGO glycated scaffold. The water sorption of the glycated scaffold did not plateau even after 50 minutes of “drying” on the ATR crystal. This indicated that the complete rehydration of glycated collagen (by H_2_O) is not a rapid process. On the contrary, the rehydration of the control collagen (non-glycated) reached a plateau after 20min. This means that our glycated collagen samples could absorb more water over a longer period of time. From these experiments, it is quite clear that the MGO glycation process impacts directly upon the water contents in collagen ^[63]^, and the interaction between collagen and the amount of absorbed water is a function of AGE formation. This outcome is supported by our original study in which we showed that collagen with riboflavin-mediated crosslinks (compared to non-crosslinked collagen) retained water for a long time upon dehydration, suggesting that crosslinks influence the collagen-water interaction.^[12]^ In another study using a combination of molecular simulations and experimental work we showed that in older tendon tissue, the presence of AGEs increases hydration and the free water content around the collagen molecules.^[23]^Previously we proposed that the increased water content in collagen resulting from glycation may act as an adaptive response to the loss of highly hydrophilic hyaluronic acid (HLA) and glycosaminoglycan (GAG) as part of the natural aging process in our skin.^[59]^ Here using a new method, we showed the superior water retention capacity of MGO glycated collagen scaffolds.

**Table 1.**
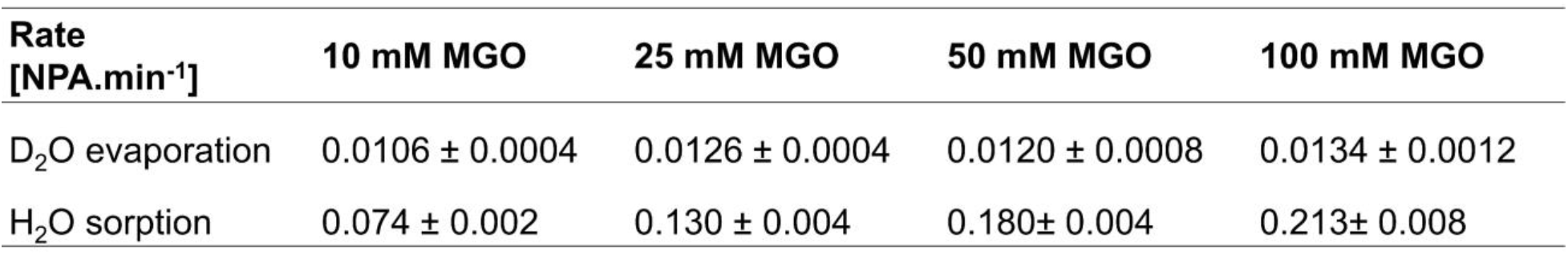
Rate of H_2_O sorption and D_2_O evaporation. Rates are calculated from the exponential term of the fitted plot in Figure 3b and presented as mean (Normalised Peak Area over time) ±SD (n=3).

| Rate<br>[NPA.min<sup>-1</sup>] | 10 mM MGO | 25 mM MGO | 50 mM MGO | 100 mM MGO |
| --- | --- | --- | --- | --- |
| D<sub>2</sub>O evaporation | 0.0106 ± 0.0004 | 0.0126 ± 0.0004 | 0.0120 ± 0.0008 | 0.0134 ± 0.0012 |
| H<sub>2</sub>O sorption | 0.074 ± 0.002 | 0.130 ± 0.004 | 0.180 ± 0.004 | 0.213 ± 0.008 |

### ● Structural and mechanical response of glycated collagen

As mentioned before, collagen molecules can self-assemble in vitro, under appropriate conditions, into microscopic fibrils and macroscopic fibers with D-banding periodicity of 67 nm that are structurally reminiscent of native collagen fibers. Atomic force microscopy (AFM) imaging was carried out on control and glycated scaffolds and single fibrils as presented in Fig. 4-a-i, iii respectively. In all samples, collagen fibrils with D-banding can be clearly observed. In the control scaffold, the orientation of the fibrils appears to be random. This is one of the characteristics of plastically compressed collagen which has been widely reported.^[64–66]^ As a function of glycation, the architectural alignment of collagen fibrils changes to become more unidirectionally oriented (Figures 4a-ii, b-ii, c-ii, d-ii). As the concentration of MGO is increased, we observed a more prominent formation of larger collagen aggregates spanning across the entire image and measuring between 10 to 20 μm long. The glycation process is promoting the formation of collagen bundles or early collagen fascicles through the interaction of neighboring fibrils sidechains with MGO-induced AGEs. These MGO-induced AGEs increase the registration of neighboring collagen fibrils leading to the formation of a homogeneous collagen sheet, which is reminiscent of collagen in the dermis. Figures 4a-iii, b-iii, c-iii, d-iii show the AFM images of control and glycated individual collagen fibrils. The characteristic D-banding is clearly visible in both control and glycated images and indicates that no significant structural or morphological change has happened after glycation at the fibrillar level. The process of glycation does not alter the morphology of the fibrils (in the dry state). Despite reports of increasing in hydrated fibril diameters as a function of glycation ^[67]^ this effect was not investigated in this study.

**Figure 4.**
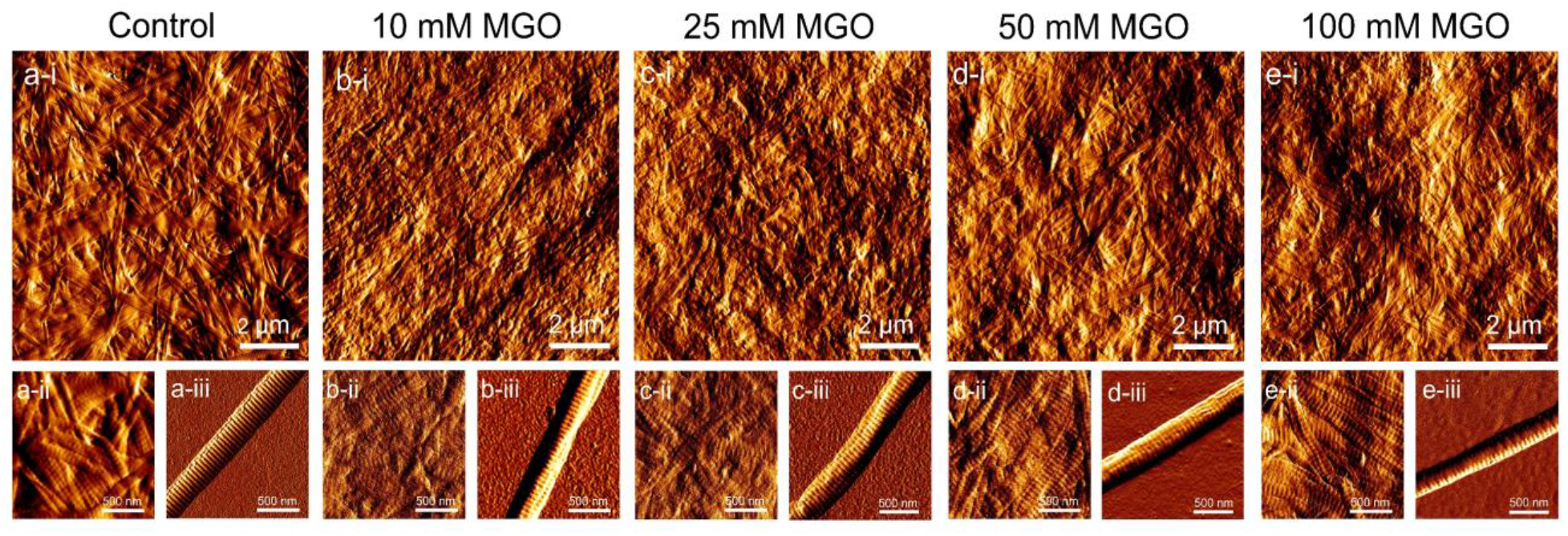
Representative AFM topological images of control and glycated collagen. recorded in ambient conditions presenting: i) the scaffold mesoscale architecture, ii) fibrillar details in the scaffold, and iii) mono-dispersed single collagen fibril. The presence of the D-banding periodicity on the engineered collagen is evident through the images and an increase in the structural alignment of collagen fibrils can be observed as a function of glycation.

For both in vivo and in tissue engineering applications, collagen is defined by its biomechanical properties in terms of load-bearing and tensile strength.^[68]^ To evaluate the impact of glycation on the mechanical properties of collagen at both the single fibrils and scaffold level, we employed AFM nanoindentation with a sharp probe (2 nm and 8 nm nominal tip radius for single fibrils) whilst ensuring that all indentations were performed on the overlap region of the collagen fibril’s D-banding periodicity and the deformations were purely elastic with no permanent deformation of the samples. Figure 5a and Figure 5b show the compressive elastic modulus of the collagen (as a function of glycation) for individual collagen fibrils (within a scaffold) and for single (isolated) collagen fibrils (not formed in a scaffold).

**Figure 5.**
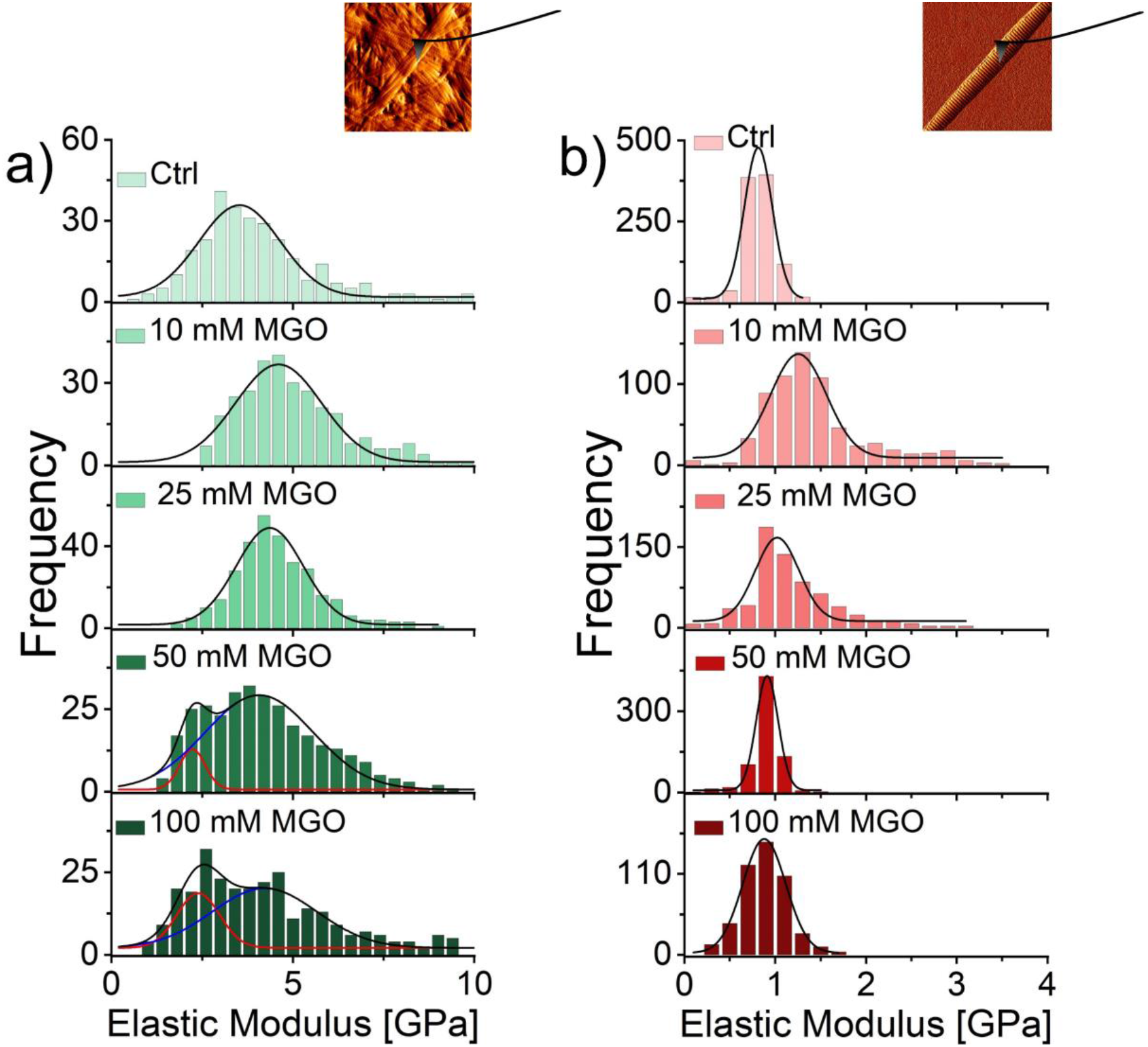
AFM nanomechanical characterization of control and glycated collagen fibrils. Frequency distributions of compressive elastic modulus of control and glycated collagen fibrils in a) the scaffold and b) mono-dispersed. The distributions were fitted (R^2^>0.95) with Gaussian distributions to calculate the Median of each distribution. Insets demonstrate the location of the indentation measurements on typical collagen fibrils. All distributions were significantly different (p<0.05) except for the mono-dispersed collagen fibrils glycated with either 50 or 100mM MGO.

The compressive elastic modulus for (non-glycated) collagen fibrils (within a scaffold) was found to be (3.53±0.06 GPa; N=300). This value is in good agreement with earlier results reported in the literature.^[69–71]^ Upon glycation (10mM MGO), the compressive elastic modulus of fibrils within the scaffold started increasing respectively to 4.60±0.06 GPa, but as the concentration of MGO increased, a bimodal distribution of elastic moduli became apparent. Interestingly, the median values of these distributions decrease as the MGO concentration increased. This bimodal distribution results from the presence of hydrophilic AGEs in the collagen fibrils. Glycation is the reaction of carbonyl groups from reducing sugars with free amino groups of lipids and proteins to form a Schiff base, which itself undergoes a time-dependent rearrangement to form a stable Amadori product. For this product to form, several conditions must be met, including the co-location of arginine and lysine residues in a confined space ^[72]^ and the diffusion of a reduced sugar at that location. This implies that collagen fibrils must experience interstitial fluid exchange to ensure the intra-fibrillar diffusion of the reduced sugars. From a structural point of view, the formation of the stable Amadori product requires the lysine and arginine residues (for example) from neighboring molecules to come in very close contact with one another. As the initial Schiff base undergoes the time-dependent re-arrangement, the distance between the original residues also changes due to the complexity and structure of the Amadori product formed.^[72]^ On the other hand, as the Amadori product is formed, it becomes hydrophilic leading to the soft binding of interstitial water as previously demonstrated. This now increases the intermolecular distance between the residues even further.^[23, 63, 72–74]^ This means that the glycation process induces first an increase in the density of the collagen fibril (Schiff base undergoing the time-dependent re-arrangement) followed by a decrease in the fibril density (hydrophilicity of the Amadori product formed). This variation in the density of the fibrils was confirmed by a mechanical indentation on individual collagen fibrils. As such the compressive elastic moduli of collagen fibrils decreased (i.e. decrease in density) as the concentration of MGO glycation increased as presented in Table 2. This trend of increase and decrease in elastic moduli was also found in the case of glycated isolated collagen fibrils (Table 2). The compressive elastic modulus of (non-glycated) isolated collagen fibrils was found to be (0.81±0.01 GPa; N=982) which is remarkably lower than the elastic modulus for (non-glycated) collagen fibrils (within a scaffold) which was found to be (3.53±0.06 GPa; N=300). This notable difference arises from the different sources and protocols used to engineer the collagen itself between these two samples as both values have already been confirmed elsewhere.í^[27]^ The variation in the elastic moduli recorded on the collagen fibrils was confirmed to be statistically significant as a function of the glycation for all groups using a Mood’s median test (p<0.05) except for the mono-disperse collagen fibrils glycated with either 50 and 100mM MGO.

**Table 2.** Recapitulative table of elastic modulus values for control and glycated collagen scaffolds. Values are presented as Median ± SD (n=300) and were obtained by fitting the frequency distribution presented in Figures 5 & 6.

| Collagen scaffold | Fibrils (In a scaffold)<br>[Gpa] |  | Single Fibrils<br>[Gpa] | Bulk (Dried)<br>[Mpa] | Bulk (Hydrated)<br>[kpa] |
| --- | --- | --- | --- | --- | --- |
| Control | 3.53±0.06 |  | 0.81±0.01 | 338.44±0.07 | 75.30±1.63 |
| 10 mM MGO | 4.60±0.06 |  | 1.25±0.02 | 291.33±0.01 | 113.14±3.47 |
| 25 mM MGO | 4.36±0.04 |  | 1.02±0.03 | 391.36±0.02 | 130.63±1.70 |
| 50 mM MGO | Peak<sub>1</sub>:<br>2.23±0.08 | Peak<sub>2</sub>:<br>4.07±0.11 | 0.91±0.00 | 593.25±0.01 | 131.77±3.03 |
| 100 mM MGO | Peak<sub>1</sub>:<br>2.39±0.13 | Peak<sub>2</sub>:<br>4.23±0.59 | 0.88±0.01 | 651.44±0.02 | 151.66±2.39 |

### ● Mechanical response of glycated collagen as a function of (re)hydration

To evaluate the effect of hydration on the mechanical properties of glycated collagen, two sets of measurements were performed: first, in an air-dried environment, then after 10 minutes of rehydration in PBS. Here, we used a large spherical probe (micrometer-size spherical tip; r=2 μm) to measure the elastic modulus of the glycated collagen scaffold as a function of hydration. AFM measurements have shown that level of hydration changes collagen mechanical properties (Figure 6).^[75–76]^ Like others, we recorded a very large decrease (5 orders of magnitude) in the elastic moduli of the control scaffolds upon hydration (E_s-dry_= 338.44±0.07 MPa and E_s-wet_= 75.30±1.63 kPa). This remarkable decrease in the elastic modulus is once again associated with a change in the density of the collagen scaffold as previously demonstrated. Upon glycation of the scaffolds, the elastic moduli of both the hydrated and dried samples follow the same increasing trend as shown in Table 2. This is an expected outcome as it is widely accepted that the process of glycation increases the stiffening of collagen. Yet, we found that the elastic moduli found for the collagen scaffolds (despite increasing) are lower than the value of elastic moduli of its constituent collagen fibrils.

**Figure 6.**
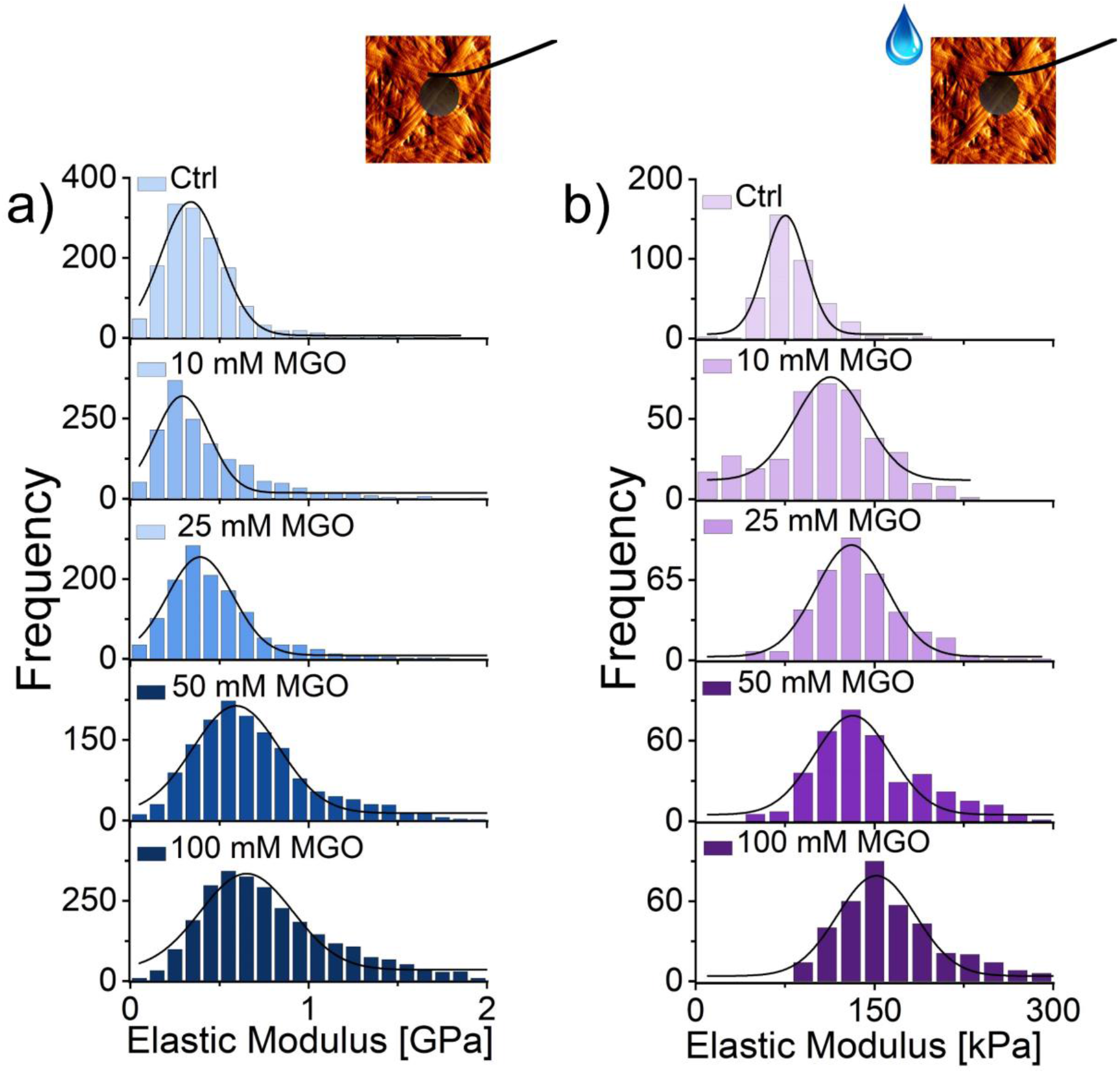
AFM nanomechanical characterization of control and glycated collagen scaffold. Frequency distributions of compressive elastic modulus of control and glycated collagen scaffold in both a) ambient and b) hydrated conditions. The distributions were fitted (R^2^>0.95) with Gaussian distributions to calculate the Median of each distribution. Insets demonstrate the location of the indentation measurements on typical collagen scaffolds. All distributions were significantly different (p<0.05) except for the ambient control and 10 mM MGO collagen scaffolds and hydrated collagen scaffold glycated with either 25 or 50mM MGO.

To understand this, one ought to review the size of the indenter in relation to the mechanical measurement performed. The indentation of the collagen using a sharp or beaded tip measures different mechanical properties of the same samples. In the case of a sharp tip (tip diameter <<< fibril diameter), the indentation cycle penetrates only a few nanometers into the fibrils (rule of 10%).^[77]^ This means that during that indentation, the compressive elastic modulus of the fibrils is measured. If the tip size is increased (tip diameter >>> fibril diameter), then the indentation cycle penetrates the scaffold rather than individual fibrils to measure the compressive elastic modulus of the scaffold.

Establishing a relationship between the compressive elastic moduli of the fibrils and of the scaffold is non-trivial. However, it is safe to assume that upon the indentation cycle using a larger beaded cantilever, the collagen fibrils in the immediate contact area between the bead and sample are being subjected to both a compressive load and a tensile load as presented in Figure 7. Thus, the scaffold compressive modulus is made of at least two mechanical components associated with the tensile and compressive elastic moduli of fibrils ^[78]^ and a poroelastic component ^[79]^ associated with reversible capillary effects. Since the compressive elastic modulus of the fibrils decreases as a function of glycation, we can then conclude that it is the composite tensile elastic modulus of the collagen fibrils that is predominantly responsible for the increase in the compressive modulus of the scaffold. There are numerous reports confirming the increase in tissue stiffening associated with the reactions between glucose and collagen which leads to the formation of advanced glycation end-products as the one described here.^[80–81]^ The macroscopic definition of elastic modulus as an intrinsic material property that is independent of size sometimes fails at nanoscale or microscale dimensions, where the moduli of materials might not only differ but also can become size-dependent ^[82–83]^ as observed in this study.

**Figure 7.**
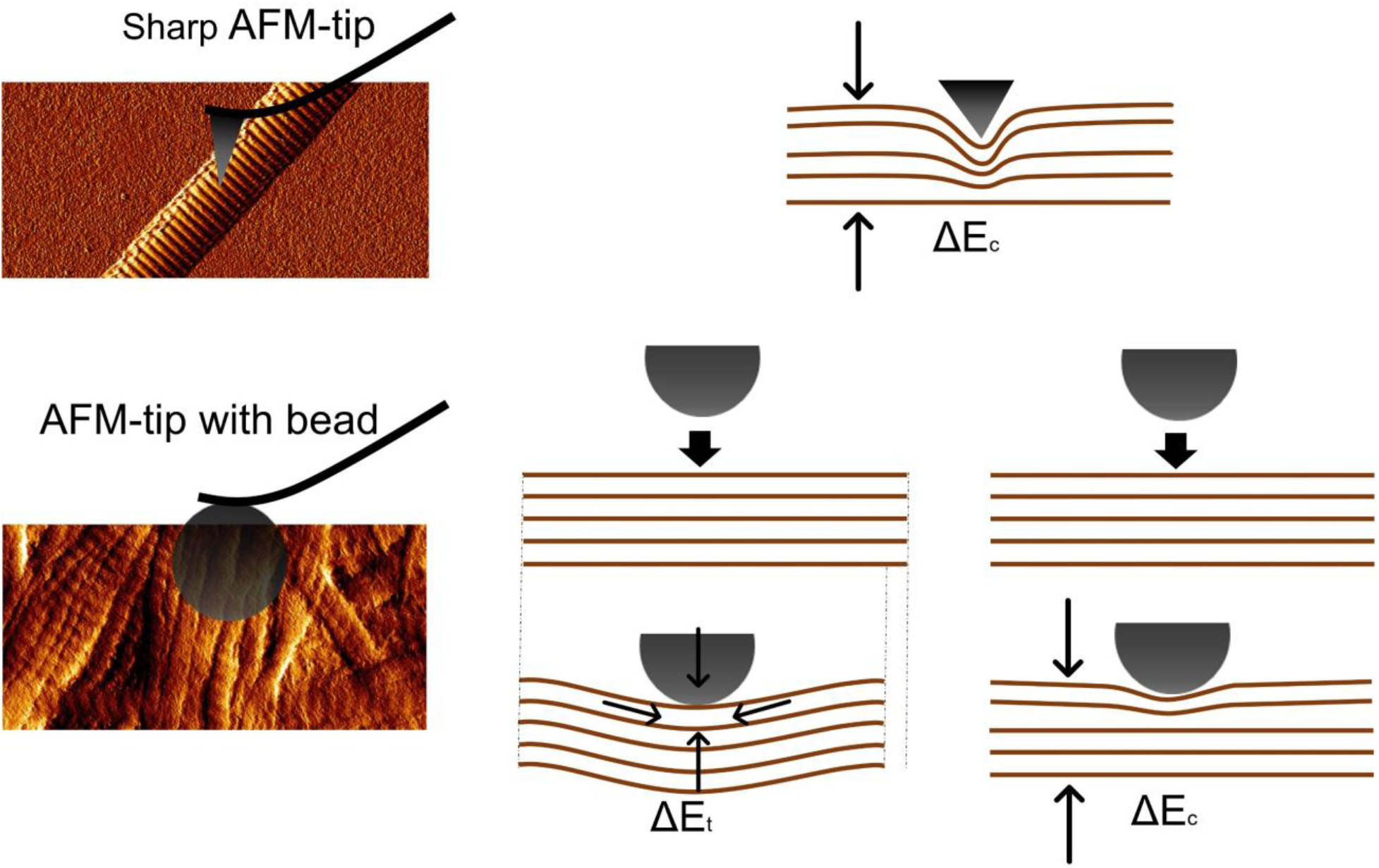
Explanatory schematic of the fibrils and scaffold elastic responses as measured by AFM indentation. The interaction of a) a nanometer-size and b) a micrometer-sized spherical AFM cantilever tip with collagen scaffold surface. A nanometer-sized tip is used to apply a compressive force to the contact surface, but using a micro-sized beaded cantilever, the collagen fibrils are being subjected to both a compressive load and a tensile load. ΔE_c_ denotes the composite compressive modulus exerted on a single fibril, whereas ΔE_t_ denotes the composite tensile modulus exerted on a single fibril. Arrows represent the direction of the force exerted on the sample.

### ● Enzymatic degradation susceptibility

Collagen is resistant to nonspecific proteinases and is susceptible to only a small number of specific collagenolytic enzymes. Bacterial collagenases and vertebrate collagenases (MMPs) are two main proteinases capable of degrading collagen. Bacterial collagenase unlike MMPs makes multiple scissions along the collagen α-chains leading to the generation of multiple small fragments.^[84]^ Defining the mechanism of collagen enzymatic degradation is important for understanding the physiological processes (e.g., wound repair and remodeling) and pathological conditions (e.g., scarring and metastasis) affecting the ECM.^[85]^ Collagen enzymatic degradation depends on different factors such as collagen intermolecular crosslinks, concentration, and type of collagenase, and collagenase solution physiochemical factors like pH and temperature.^[86]^ The enzymatic degradability of the collagen scaffolds was evaluated by a bacterial collagenase (Clostridium Histolyticum Collagenase (CHC)) and the effect of crosslinking on the rate of degradation was studied at 37 °C until complete degradation was observed by timelapse digital imaging. Visual inspection revealed that glycated scaffolds by 100 mM MGO exhibited the highest resistance to enzymatic degradation, while non-glycated scaffolds exhibited the least resistance to enzymatic degradation (Figure 8). The non-glycated scaffold was degraded entirely after 100 min exposure to collagenase. Crosslinking significantly slowed degradation in comparison to control. Complete degradation for 10 mM MGO, 25 mM and 50 mM MGO collagen scaffolds was observed after 100 min, 160 min, and 450 min, respectively. For 100 mM MGO scaffold, complete degradation took approximately 3 days.

**Figure 8.**
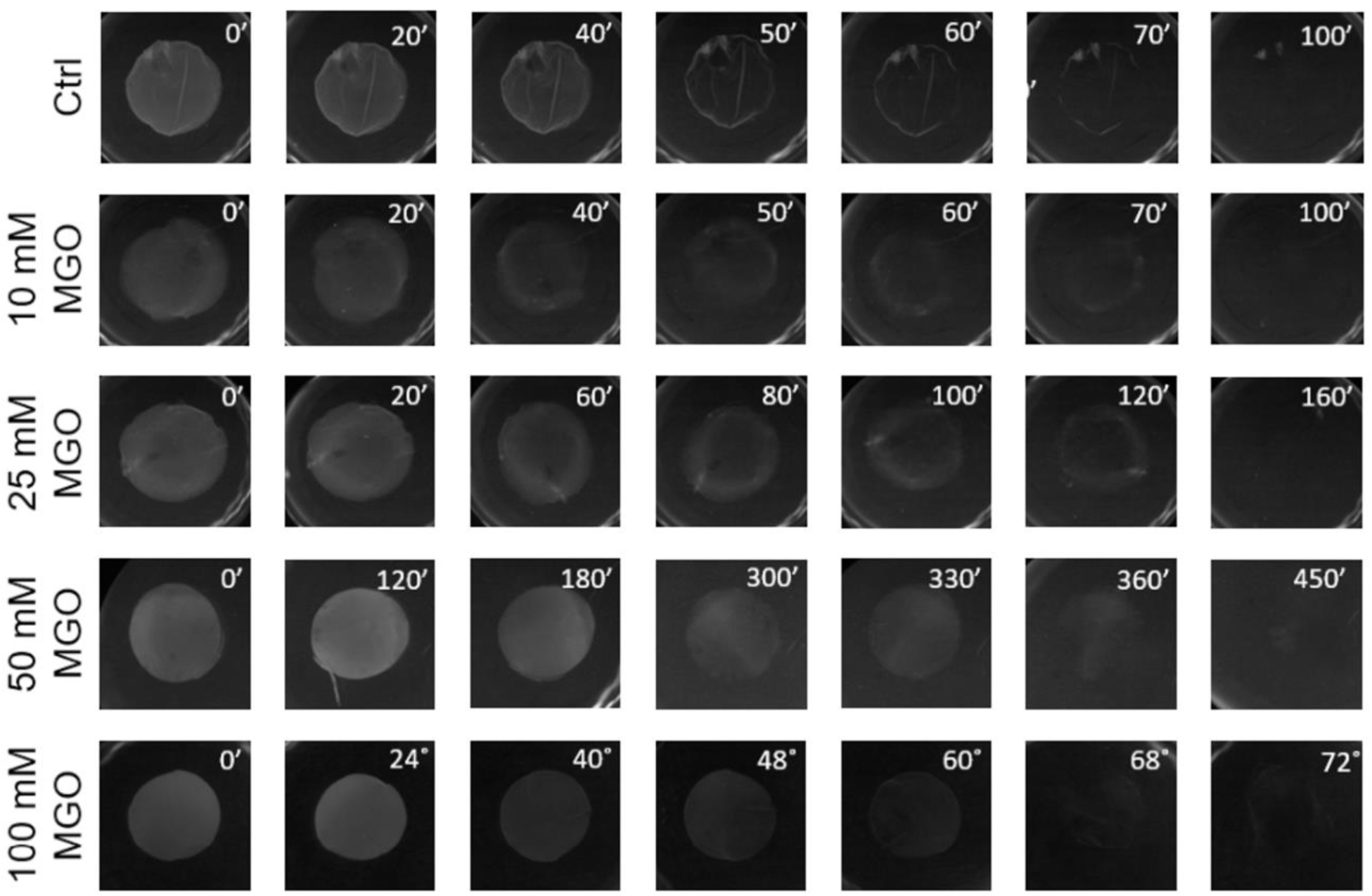
Time-lapse digital imaging (7X) of control and glycated collagen scaffold undergoing enzymatic degradation by Clostridium histolyticum collagenase (CHC). Times are denoted in min: “ and hour: °.

As demonstrated in Figure 9 collagen scaffolds are degraded over time following a sigmoidal degradation pattern. The initial slow degradation phase is followed by a rapid phase transition of the fibril (gelatinization – fibrils fragmentation). We can therefore assert that the enzymatic degradation is not a linear degradation process and that the tissue (here collagen) does both mechanically and structurally fail before the complete digestion of the matrix. The phase transition observed in the sigmoidal curve is reminiscent of the pattern of degradation of collagen under heat denaturation ^[87]^ in which the collagen suddenly denatures (gelatinizes) at 65°C following an early onset of degradation at 58°C. The slope of the phase transition and onset of degradation (start point of rapid degradation) can be used to compare the rate of degradation in different samples which are reported in Table 3. The onset of degradation measured here increased significantly as a function of MGO exposure reaching up to 31.7 hrs for the 100 mM MGO group. This is an increase of over 85x in terms of resistance to enzymatic degradation for the collagen scaffold. The phase transition rate also demonstrates the increased resistance to degradation of the glycated collagen as the rate measured during the phase transition decreased by 2 orders of magnitude as the glycation increases. We can therefore conclude that glycation not only slows down the initiation of collagen degradation, but it also slows down the rate at which it occurs. To understand this, it is important to consider the role of the MGO-induced AGEs in collagen. As discussed previously, these crosslinks bind two adjacent collagen molecules covalently. As the bacterial collagenase attacks the collagen fibrils, it non-specifically cleaves the superficial collagen molecules leading to the loosening and collapsing of the fibrillar structure. This is evidenced in the AFM topology images of the scaffolds recorded post-degradation as presented in Figure 10. Other studies have also demonstrated a decrease in collagen degradation rates as a result of crosslinking.^[85, 88]^ It has been shown that several residues involved in AGE crosslinks are binding sites for collagenase which prevents collagenase from engaging, resulting in decreased collagen affinity for collagenase.^[86, 89]^ The presence of MGO-induced AGE crosslinks partially inhibits this degradation mechanism by preventing the loosening of the fibrillar morphology by providing mechanical reinforcement between the attacked collagen molecules themselves. This suggests that the accumulation of MGO-induced AGEs crosslinks acts as an internal scaffold within the fibril supporting the fragmented surface collagen molecules. However, the fibril eventually collapses when too many collagen molecules have been cleaved. AFM images of degraded scaffolds show a clear decrease in fibril diameter and slowly fading of the D-banding over time. Before degradation, there are intact fibrils with clear D-banding and through degradation, a reduction in fibril diameter is observed. At the final stages of degradation, there is no evidence of banding periodicity, just an amorphous mass on the surface with remaining fragments of collagen fibrils (images not shown). In our approach, we demonstrated that we could modulate the rate of degradation with in vitro glycation which will be of service for tissue engineering and clinical applications in the future ^[90–91]^. It is worth mentioning that the complex in vivo degradation due to cells and other enzymes cannot be entirely replicated through in vitro enzymatic degradation assay.

**Figure 9.**
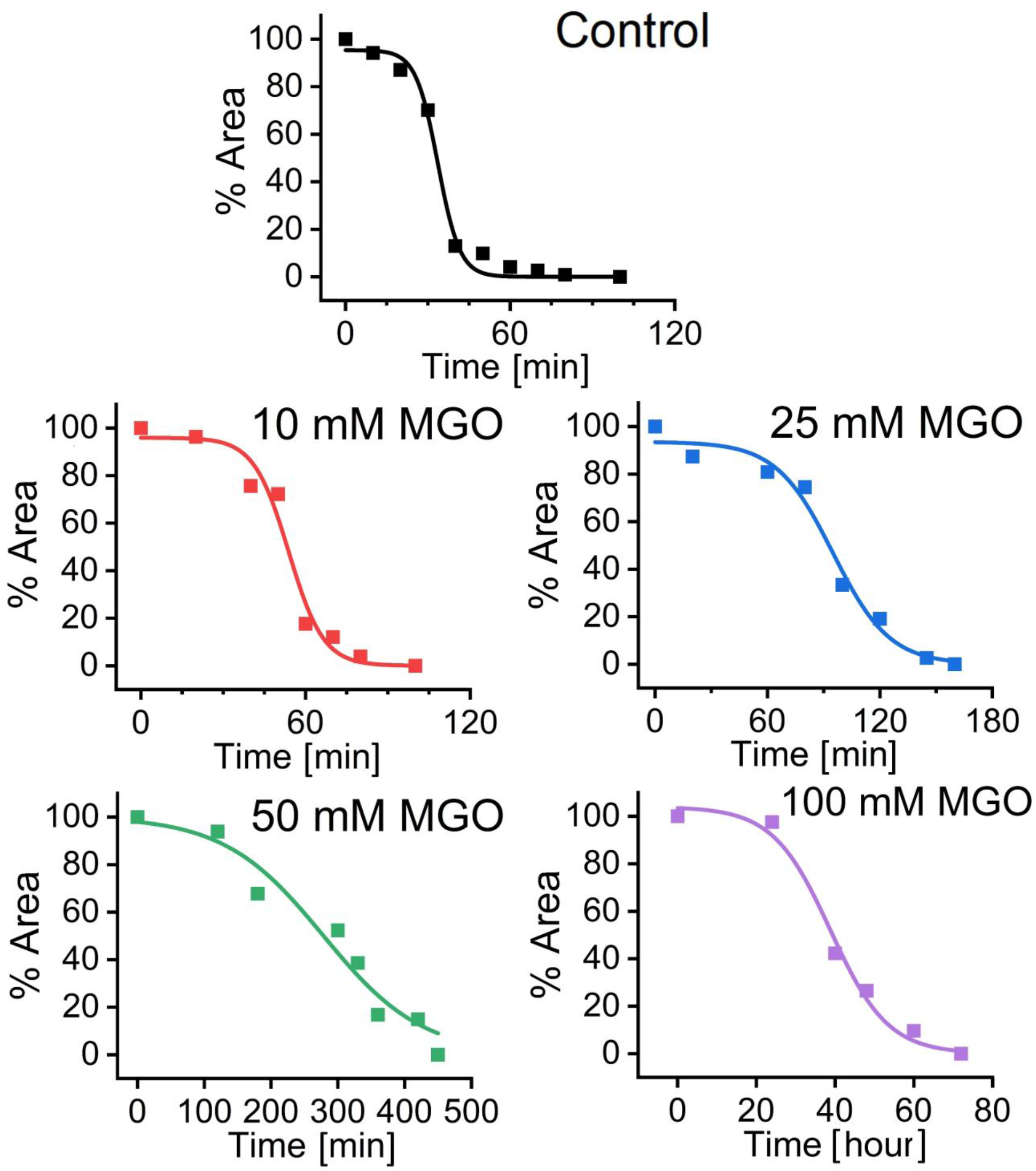
Enzymatic degradation susceptibility. Representative collagen scaffolds enzymatic degradation plots obtained by measuring the reduction in the scaffold surface area as a function of time. The plots were fitted with a sigmoidal curve (R^2^>0.95).

**Figure 10.**
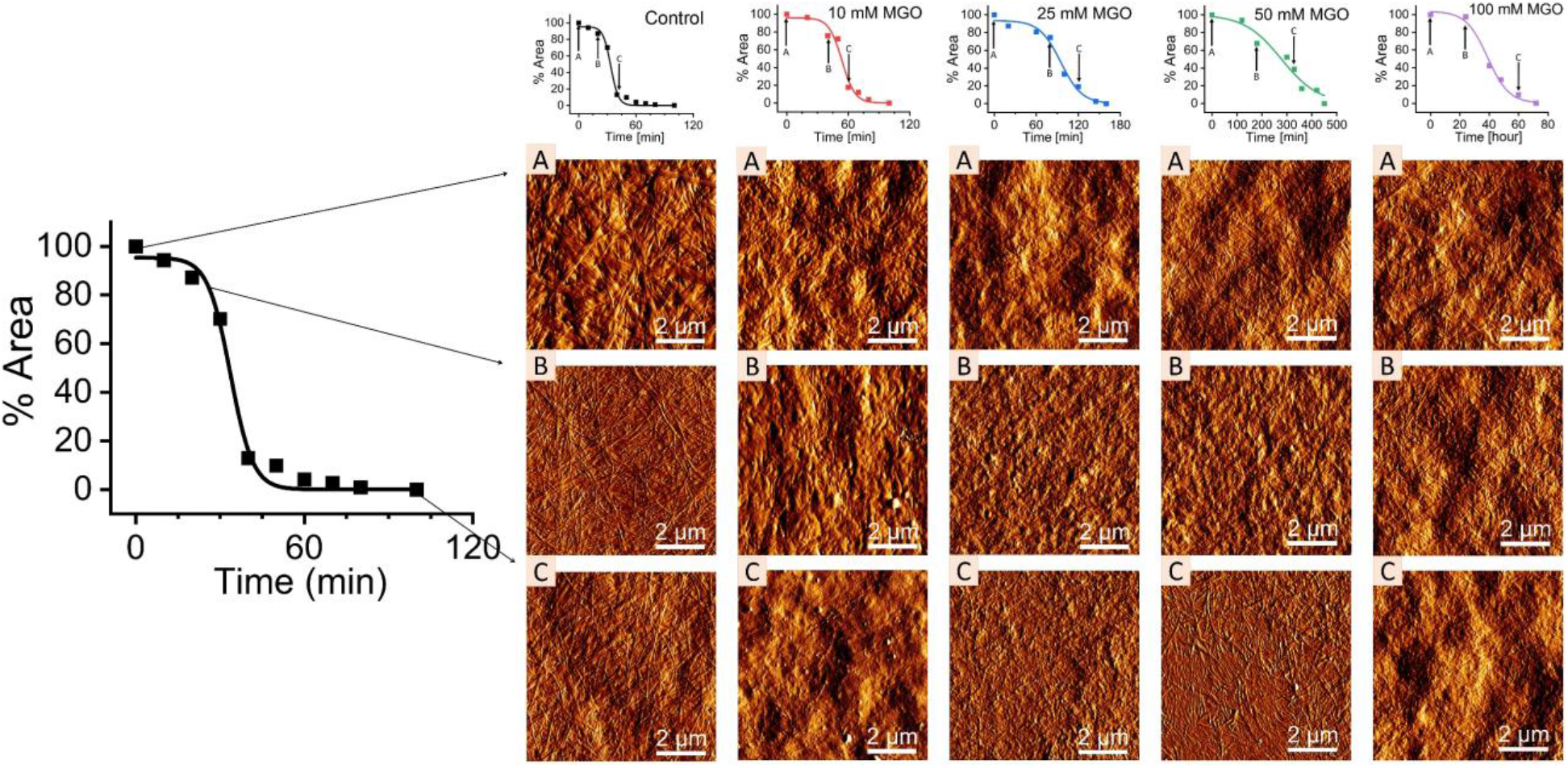
Representative AFM topological images of control and glycated collagen scaffolds obtained at specific time points of A) start of degradation and post-degradation B) the onset of degradation and C) almost completed degradation.

**Table 3.** Degradation rate of collagen scaffolds by bacterial collagenase (CHC). Phase transition rates are presented as Median ± SD (n=3) and were obtained by fitting the degradation plots presented in Figure 9.

| Collagen scaffold | Onset of degradation | Phase transition rate<br>[$\delta\%$ Area/min] | Time of complete<br>degradation |
| --- | --- | --- | --- |
| Control | 21.75 min | $0.25 \pm 0.04$ | 100 min |
| 10 mM MGO | 46.29 min | $0.16 \pm 0.04$ | 100 min |
| 25 mM MGO | 91.86 min | $0.07 \pm 0.01$ | 160 min |
| 50 mM MGO | 277.47 min | $0.01 \pm 0.00$ | 450 min |
| 100 mM MGO | 31.7 hour | $0.002 \pm 0.00$ | ~ 3 days |

### ● Application towards Tissue Engineering

While the development of collagen scaffolds for clinical applications is a sustained area of research in tissue engineering, researchers have not yet been able to recreate a truly biomimetic tissue. Different approaches are often taken depending on whether the development is focussed on either cell-based engineering on the collagen layer or structural collagen scaffold engineering strategies. Although both strategies ought to lead to clinical applications, the collagen fibrillogenesis quality control appears different for both these approaches. In fact, few cell-based studies are concerned with the presence of D-banding periodicity on the collagen fibrils often used as a cell substrate. This is particularly important since cellular phenotypes are modulated by their environment ^[92]^ in which the collagen morphology plays an important part. Here, we observed that collagen can present unique morphologies, ranging from a D-banded fibrillar matrix to a gelatinized amorphous layer. Our quality control started by ensuring that our collagen was indeed in the form of a D-banded fibrillar matrix. In our study, we used atomic force microscopy to confirm this but other routine approaches such as a Picro-Sirius Red staining combined with polarisation microscopy could also confirm the presence of these bands.^[93]^ The early formation of collagen fibril bundles or early fascicles has not yet been demonstrated in vitro unless the scaffold was forcibly aligned ^[94] [95]^ during the fibrillogenesis stage. Combining this ongoing effort for achieving a high level of collagen alignment with a crosslinking method promoting fibrils bundle formation would offer a realistic method to engineer biomimetic ligaments and tendons. A second limitation related to the engineering of collagen scaffolds is the reduced biomechanical properties of the scaffold created in relation to the tissue to be modeled or repaired.^[96]^ This has been a long-standing problem in tissue engineering and various solutions have been brought forward to crosslink collagen. While the recent review by Adamiak ^[97]^ discusses a more recent approach to collagen cross-linking using glutaraldehyde, genipin, EDC-NHS, dialdehyde starch, chitosan, temperature, UV light, and enzymes, the in vitro endeavor to increase the mechanical properties of collagen dates back many decades as exemplified by early work from Siegel who used highly purified Lysyl Oxidase to crosslink chick bone collagen fibrils.^[98]^ The literature accounts for three types of polymerization techniques that can reinforce the collagen structure: physical, chemical, and enzymatic cross-linking, but is omitting to include glycation as a potential mechanism for reinforcing collagen. Yet, the use and formation of advanced glycation end-products crosslinks are routinely used in the case of clinical treatment of keratoconus— the most common degenerative dystrophy where the cornea loses mechanical stability. In the work presented by Mi et al.^[99]^, riboflavin crosslinking has been used in the treatment of progressive keratoconus.^[100–101]^In our approach, we demonstrated how we can harness the rapid and highly efficient collagen crosslinking potential of MGO. The concentrations used in our approach exceeded by several orders of magnitude the native concentration found in an aging individual, yet we engineered crosslinks that natively exist in the human body. Using this approach, we managed to modulate the glycation mechanism to confer heightened mechanical and proteolytic degradation resistance to collagen scaffolds that are structurally reminiscent of native tissues. Through this approach, we also overcome one of the most challenging issues related to the tissue engineering of collagen scaffolds, namely the rapid matrix turnover. Using our crosslinking approach, we increased by a factor of 85x the resistance to enzymatic degradation of our collagen scaffolds. This has a tremendous application towards collagen-based membranes ^[102]^ for guided bone regeneration (GBR) in both dentistry and orthopedics.^[103–104]^ Despite their widespread use in clinical medicine and dentistry, commercially available guided bone regeneration membranes present significant drawbacks that can affect the new bone formation resulting in delayed or compromised healing. These drawbacks are well-known amongst practitioners and include weak mechanical strength, rapid degradation, or non-degradation of the membrane. The GBR membrane’s biocompatibility has been largely achieved by using collagen as a base scaffold. However, the tuning of the turn-over of these collagen-based membranes still poses a significant unmet clinical challenge that can be overcome through controlled glycation as presented here. Finally, the major advantage of a controlled glycation approach to enhance the properties of collagen scaffolds resides in the increased affinity of the glycated collagen with water. In vivo, collagen is mostly co-localized with water-binding glycans. The affinity between collagen and water is essential for matrix homeostasis and is related to the abundance and location of these glycans. While the engineering of collagen-glycans is possible, it is harder to modulate the physical properties of the collagen in the presence of such glycans as these molecules tend to surround the entire collagen fibrils. Thus, our approach once more proves promising in binding water within the collagen fibrils especially essential for ligaments engineering.

## Conclusion

For the last three decades, collagen has been used, engineered, and modified to create new scaffolds for tissue engineering and clinical applications. As the fundamental mechanisms governing the formation and remodeling of collagen both in vitro and in vivo are being uncovered, new approaches are being developed to harness these mechanisms to engineer these collagen scaffolds with increased biomimeticity. Harnessing glycation is a challenge as it is often associated with pathological conditions in humans. Yet, the crosslinking chemistry of AGEs offers many advantages towards tissue engineering compared to more synthetic approaches. Here, we showed how we could use a rapid-reacting reduced sugar to modify both collagen scaffolds and fibrils by conferring unique biophysical and biochemical properties. Understanding the mechanism of glycation on the biophysical and biochemical properties of collagen was made possible only because our starting collagen scaffolds did not possess any covalent intermolecular crosslinks. The properties evaluated were: well-defined collagen ultrastructure at both fibril and scaffold levels, formation of fibril bundles, increased scaffold stiffness without increasing the individual fibril stiffness, the increased affinity of collagen to water, and finally modulation of the collagen resistance to proteolytic degradation. While our study considered only one source of AGEs, reduced sugars are numerous as they accumulate in our bodies as we age. This study demonstrates how we can redeploy our understanding of collagen biophysical and biochemical properties towards the in vitro design of collagen scaffolds with biomimetic properties.

## Author Contributions

MV, MA, and LB contributed to the conception, design, data acquisition, analysis & interpretation and drafted the manuscript, LH, SH, GB and KG contributed to the acquisition, analysis & interpretation of the data. Finally, SA and CS contributed to the interpretation of the data. All authors critically revised the manuscript before submission. All authors gave their final approvals and agreed to be accountable for all aspects of the work.

## Materials and Methods

### Engineering collagen fibrils and scaffolds

Mono-dispersed collagen fibrils were produced by mixing 4 μl of type I tropocollagen monomers (3mg/mL in 0.01 N HCl, pH 2) from human placenta in acidic solutions (Advanced BioMatrix Inc., Carlsbad, CA) with a solution composed of 6 μl of UHQ water, 20 μl of 200 mM Na_2_HPO_4_ adjusted to pH 7 with HCl, and 10 μl of 400 mM KCl. The neutralized solution was then placed on a shaker at 37 °C for 6h to enable fibrillogenesis. The solution of mono-dispersed collagen fibrils was then stored in an incubator at 37 °C until use (<7 days).^[27]^

Collagen scaffolds were prepared by adding 3ml rat-tail tendon collagen type I solution (2.0 mg/ml protein in 0.6% acetic acid; First Link Ltd, UK) to 1ml 10× Eagle’s minimum essential medium (MEM). The solution was neutralized using 5M and 1M NaOH, transferred to a 24-well plate (1 ml in each well), and then placed in a humidified incubator at 37 °C for 1 hour to allow fibrillogenesis.^[12, 28]^ The collagen hydrogels were then plastically compressed to increase their density as performed elsewhere (Figure 1a).^[29]^ Following compression, the scaffolds were stored in PBS at 4 °C until use (<7 days).

### Collagen glycation by Methylglyoxal

Both collagen fibrils in solution and collagen scaffolds were crosslinked using Methylglyoxal (MGO) (Sigma-Aldrich) post-fibrillogenesis. Serial dilutions of MGO were used to create 10 mM, 25 mM, 50 mM, and 100 mM MGO solutions (from a stock 40 % aqueous solution) in PBS (adjusting each solution to pH 7.4). First, collagen fibrils in solution were physisorbed onto a glass substrate for 2hrs before the excess collagen solution was rinsed off using a laminar flow of UHQ water. The glass coverslips with the physisorbed collagen fibrils were then placed inside Petri dishes before being immersed in the MGO solutions (at defined concentrations) and incubated for 3 days at 37 °C. Following their plastic compression, the collagen scaffolds were transferred into Eppendorf tubes containing the different concentrations of MGO solutions and incubated for 3 days at 37 °C. After 3 days of incubation, all collagen samples were rinsed using a laminar flow of UHQ water and stored into PBS at 4°C until characterization.

### Autofluorescence

MGO-derived AGEs formation was determined by assessing the autofluorescence of the glycated collagen scaffolds. For this purpose, 50 μl of newly formed collagen scaffolds were incubated with 100 μl of MGO solutions at various concentrations in a black 96-well plate (Nunc, Rochester, NY). The fluorescence intensity of crosslinked collagen was measured at 420 nm emission/340nm excitation using a multimode microplate reader (Cytation 3, BioTek, USA).^[17]^ The fluorescence intensity for each crosslinked collagen was measured in triplicate at 24-h intervals over 3 days before being averaged and then compared to the control.

### Fluorescence Lifetime Imaging(FLIM)

Time-correlated single photon counting (TCSPC) FLIM was performed with an inverted confocal laser scanning microscope (A1R, Nikon, Japan) with a FLIM add-on module (LSM Upgrade Kit, PicoQuant, Germany) and SymPhoTime 64 software for data acquisition and fitting (PicoQuant, Germany). Fluorescence was excited with a picosecond pulsed 405 nm laser operating at a 5 MHz repetition rate, and the photons from the decay were collected with a hybrid PMT over a period of 200 ns with 50 ps bin width. To avoid photon pile-up, excitation intensity was adjusted such that the data collection rate did not exceed 1% of the excitation repetition rate. The samples were placed on a coverslip and imaged with a 20X NA0.75 objective (Nikon, Japan). A randomly chosen area of 213×213 μm was scanned with 512×512 pixels until the peak of the sum decay had at least 10^6^ photons, and decays from each pixel were added together for lifetime analysis. The experiment was repeated for a total of three randomly chosen areas from each sample, and two samples for each MGO concentration.

The sum decay for each area was fitted with a three-exponential function:

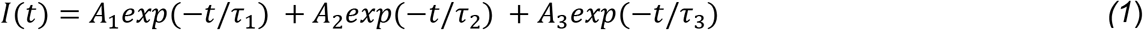

where *I*(*t*) is the measured fluorescence intensity, *τ*_1_, *τ*_2_ and *τ*_3_ are the decay times, and *A*_1_, *A*_2_ and *A_3_* are their corresponding amplitudes at *t* = 0.

The mean fluorescence lifetimes *τ_A_* and *τ_B_* were determined from equation (2):

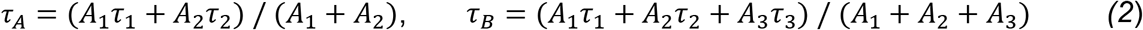

where *τ*_1_ and *τ*_2_ correspond to the two shorter lifetime components from equation (1), and *A*_1_ and *A_2_* are their corresponding amplitudes; and *τ*_B_ includes all decay components. The average lifetimes *τ_A_* and *τ_B_* were then calculated for each MGO concentration from the six measurements per concentration, and errors obtained from the standard deviation.

### AGEs Competitive ELISA Assay

The formation of MGO-derived AGEs was also measured by using a competitive ELISA kit (ab238543, Abcam) according to the manufacturer’s instructions. Briefly, protein binding 96-well plate wells were coated with 100 μl MGO conjugate in PBS (500 ng/ml) and incubated overnight at 4°C. Wells were washed with 1X PBS and blocked with 200 μl assay diluent for 1 hour at room temperature. For sample preparation, collagen scaffolds were snap-frozen in liquid nitrogen and pulverized before being transferred into a 1.5 ml Eppendorf tube. 100 μl RIPA buffer was added to the samples, followed by sonication and centrifugation. Total protein content was measured using BCA assay, and samples were adjusted to 100μg/ml in RIPA buffer. The coated wells were incubated with a dilution series of standards or samples (50 μl) for 10 min, and then 50μl 1x anti-MGO antibody was added and incubated for 1 hour on an orbital shaker at room temperature. The wells were washed 3 times with wash buffer and incubated with the secondary antibody-HRP conjugate. Finally, substrate solution was added, and the reaction was stopped by stop solution. Optical density (OD) value of the samples was measured using an absorbance microplate reader (EPOCH 2, BioTek, USA) at 450 nm wavelength. Following calibration against standards, the value OD measured from the glycated collagen could be used directly for direct quantification of MGO-derived AGEs in our scaffolds. All measurements were carried out in triplicate and presented as mean +/− stdev.

### Fourier-transform infrared spectroscopy (FTIR)

Control and glycated collagen scaffolds placed on CaF_2_ salt windows were imaged with an Agilent Cary 670 spectrometer and 620 infrared microscope equipped with a 15x (0.62 NA) objective and a 64×64 Focal Plane Array mercury cadmium telluride detector. The sample was imaged through with slide-on micro-ATR accessory (Ge crystal) yielding a nominal geometric pixel resolution of 1.4 × 1.4 μm^2^. All FTIR-ATR spectrochemical images were collected as a sum of 128 sample scans, ratioed against 512 background scans recorded against air, at 4 cm^−1^ spectral resolution.

For H_2_O/D_2_O exchange experiments, control and glycated collagen scaffolds were immersed in deuterium oxide (D_2_O) for 48 hrs in Eppendorf tubes at room temperature. Upon their removal from the D_2_O, the collagen scaffolds were directly placed (while hydrated) on the diamond windows of a GladiATR (GladiATR, Pike technologies, USA) mounted inside an iS20 FTIR spectrometer (Nicolet, Thermo Scientific, USA). The ATR spectra were recorded at 5 min intervals (resolution 4cm^−1^, 32 co-additions) from the moment the scaffolds were mounted on the diamond windows until they dried up (~50min). The relative variations in the D_2_O stretch (2625cm^−1^) and the OH-stretch (3,530 cm^−1^) were used to plot the rate of D_2_O evaporation and H_2_O absorption over time as a function of the glycation of the scaffold. The evaporation and absorption plots were then fitted with a first-order exponential growth (absorption) or decay (evaporation) while ensuring that the fitting R^2^>0.9.

### Enzymatic Degradation

The enzymatic degradation susceptibility of the collagen scaffolds was evaluated by incubating the control and glycated collagen scaffolds with collagenase (*Clostridium histolyticum* (CHC), Sigma-Aldrich) at 37 °C. To do so, a 6.5 mm hole punch was used to homogenize the scaffolds’ surface area. Each sample was placed in a 24-well plate containing 1ml of Tris-HCl buffer and 0.25 mg/ml collagenase. The buffer was prepared by mixing 50 mM Tris, distilled water, and 5 mM CaCl2. The pH of the buffer was adjusted to 7.4 by adding HCl. The collagen scaffolds were digitally imaged at fixed time intervals using a stereomicroscope (Nikon, SMZ800, Canada) equipped with a 7X objective during the enzymatic digestion until complete degradation. The reduction of the surface area of the scaffolds was obtained by processing the time-lapse digital images on Image J (version 1.44) using a contour analysis routine. The reduction of the surface area for each scaffold was then plotted at a function of time before being fitted with a sigmoidal curve. Three regions were empirically defined based on the shape of the fitted sigmoidal curve – namely the onset of degradation, the phase transition, and the completed degradation. The rate of degradation was calculated during the phase transition by fitting a linear regression. All measurements were carried out in triplicate and presented as mean +/− stdev.

### Atomic force microscopy

A JPK Atomic Force Microscope (JPK Nano-wizard@4 BioScience, Bruker, Germany) was used for imaging and force spectroscopy of the collagen fibrils and scaffolds. Collagen imaging was performed in contact mode under ambient conditions using MSLN-10-C (Bruker, Germany) and SCOUT-70 cantilevers (Nu Nano, UK). 10×10 μm and 2×2 μm images were typically recorded with optimized settings at 1 to 2Hz. Collagen indentation was undertaken in both hydrated (PBS) and air-dried conditions (24h). All indentations were carried out exclusively on the D-bands (overlap region) of identified collagen fibrils when these D-bands could be observed by imaging. For these measurements on individual fibrils, FESPA-V2 cantilevers (Bruker, Mannheim, Germany) with a nominal spring constant of 2.8 N/m were used. For indentation measurements on the collagen scaffolds, FMR cantilevers (Nanotools, USA) with a spring constant of 2.8 N/m were used. For each sample (3 samples per group), a minimum of 300 indentation curves was obtained at no less than 5 locations across each sample (300 individual fibrils). The (compressive) elastic modulus E was obtained from all the measurements by fitting the Hertzian model to the force-distance curve obtained. The resultant modulus values were then plotted as a distribution to obtain the median and error for the control and glycated samples. All images were processed, and data analysis was performed using the JPK data processing software (version 6.3.5).

### Statistical analysis

Statistical significance was assessed with either One-way ANOVA, or Mood’s median tests. Significant differences were defined as p < 0.05 (indicated with an asterisk in the figures). Results were reported as means +/− standard deviations. All statistical analyses were performed with OriginPro 2021 software.

## Acknowledgments

The authors would like to thank NSERC for supporting the research under the project grant 512186 and the Faculty of Dentistry at the University of Toronto for supporting the research students financially. The authors would like to acknowledge the use of the CAMiLoD Imaging Facility at the Faculty of Dentistry, University of Toronto. The authors acknowledge the facilities, and the scientific and technical assistance of Microscopy Australia at the Centre for Microscopy, Characterisation & Analysis, The University of Western Australia, a facility funded by the University, State, and Commonwealth Governments. The authors declare no potential conflicts of interest with respect to the authorship and publication of this article.

